# Enzymatic metabolon improves kinetic efficiency of reaction-limited enzyme pathways

**DOI:** 10.1101/2023.07.17.549414

**Authors:** Srivastav Ranganathan, Junlang Liu, Eugene Shakhnovich

## Abstract

In this work we investigate how spatial proximity of enzymes belonging to the same pathway (metabolon) affects metabolic flux. Using off-lattice Langevin Dynamics (LD) simulations in tandem with a stochastic reaction-diffusion protocol and a semi-analytical reaction-diffusion model, we systematically explored how strength of protein-protein interactions, catalytic efficiency and protein-ligand interactions affect metabolic flux through the metabolon. Formation of a metabolon leads to a greater speed up for longer pathways and especially for reaction-limited enzymes while for fully optimized diffusion-limited enzymes the effect is negligible. Notably, specific cluster architectures are not a prerequisite for enhancing reaction flux. Simulations uncover the crucial role of optimal non-specific protein-ligand interactions in enhancing catalytic efficiency of a metabolon. Our theory implies and bioinformatics analysis confirms that longer catalytic pathways are enriched in less optimal enzymes while most diffusion-limited enzymes populate shorter pathways. Our findings point towards a plausible evolutionary strategy where enzymes compensate for less-than-optimal efficiency by increasing their local concentration in the clustered state.

## 1 Introduction

Metabolism is critical for sustaining life as it allows cell to carry out essential functions such as energy production, biosynthesis of molecules, and elimination of waste. Biochemical activity is carried out via metabolic reaction pathways, which are often interconnected and composed of multiple enzymes that catalyze specific chemical reactions ^1, 2^ in a long pathway. That is to say, the network of enzymes and metabolites functions collectively to fulfill each metabolic reaction pathway ^2–4^. Such complexity gives rise to interesting questions. How do the cells regulate and optimize different metabolic pathways in an efficient manner within crowded intracellular environments? Could higher-level organization of enzymes enhance the efficency of metabolic pathways? If so, what spatial organization of enzyme networks can be most effective for metabolic processes?

Spatial enzyme clustering could influence metabolism by selective segregation or sequestration of metabolic pathway components ^5^. Enzyme clustering into spatially homogenous clusters is reported to speed up the processing of intermediates, and subsequently achieve higher metabolic reaction flux ^6^ in multi-enzyme pathways. Such efficiency enhancement mainly comes from concentration of enzymes from the same metabolic reaction pathway into dense clusters. Enzyme clustering can also benefit reaction through increasing local substrate concentration, the reason behind which can be well explained by Michaelis-Menten kinetics. Thirdly, enzyme clustering may alter reaction kinetics via a change in local environment, such as pH, ionic strength^7^ which modify the intrinsic activity of those enzymes. Moreover, local molecular conformation changes can also lead to the increase/decrease in an enzyme’s catalytic efficiency ^8^. Besides enzyme clustering, there are also channeling mechanisms, including direct and proximity channeling, although their biological implications are still widely debated ^9, 10^.

Most models developed so far focus on short metabolic reaction pathways ^6^, which are less biologically relevant. How enzyme clustering facilitates longer metabolic reaction pathways as opposed to single enzyme kinetics remains unknown. Previously, Castellana et al.^6^ have established how density modulation due to clustering of enzymes improves the rate of intermediate processing and speeds up reactions. However, their seminal semi-analytical work does not systematically explore how clustering would influence enzymes that vary in their enzyme catalytic efficiency (k_cat_/K_m_). Further, how the nature of the physical interactions between substrates and enzymes could tune catalytic efficiency has also not been studied in the context of enzyme clusters.

In this context, distribution of *k*_*cat*_/*K*_*M*_ values of prokaryotic and eukaryotic enzymes shows that only a small fraction of all enzymes fall within the diffusion-limited^11^ regime. This surprising observation that most enzymes are four-five orders of magnitude slower than the “perfect” diffusion-limited enzyme suggests that there must be other potentially generic mechanisms at play to improve efficiency of enzymatic pathways. Could enzyme clustering serve such a role to improve the efficiency of otherwise imperfect enzymes that operate far from the diffusion-limit? How does pathway length influence efficiency of reactions in enzyme clusters? How does the nature of protein-ligand interactions outside of the binding pocket influence reaction efficiency in enzyme clusters?

In this paper, we first address these key questions using off-lattice Langevin Dynamics (LD) simulations coupled with a stochastic reaction-diffusion protocol. The LD simulations capture the speedup in reaction flux in the regime of protein-protein interactions that favor clustering of enzymes. For the first time, we systematically explore the parameter space varying parameters such as protein-protein interaction strength, reaction rates (reaction probability) and non-specific protein-ligand interaction strength. Our simulations reveal that clustering is most effective in speeding up metabolic flux for reaction-limited enzymes while for fully optimized diffusion-limited enzymes the gain from clustering is insignificant. Further, rather counterintuitively, we show that there exists an optimal non-specific protein-ligand interaction strength that results in a maximal speedup in the clustered phase. We further test the findings from our LD simulations using a computationally tractable, semi-analytical approach previously proposed by Castellana et al ^6^. The semi-analytical approach allows us to sample a wider range of catalytic efficiencies and pathway lengths and further confirms the finding that long reaction pathways that involve imperfect, reaction limited enzymes show the highest speedup upon clustering. The key implication from our simulations and semi-analytical theory is that longer metabolic pathways are enriched in less optimal reaction-limited enzymes while highly optimized diffusion-limited enzymes populate shorter metabolic pathways. Bioinformatics analysis confirms this prediction showing statistically significant relation between catalytic efficiency of enzymes and the average pathway lengths they are part of.

## Methods

### Langevin Dynamics (LD) Simulations

#### The Patchy Particle Enzyme Model

Hard-sphere patchy-particle are commonly employed to simulate self-assembly of multivalent proteins ^12–14^. Here, we model enzymes as hard spheres with an adhesive interaction site that represents the active site of the enzyme (Fig. 1A). Each active site patch has a specific identity that is complementary to a unique type of ligand particle. Ligand (substrate) molecules are represented by hard-spheres that diffuse in the simulation box and react upon binding to the complementary active site. In Fig.1A we present a schematic representation of the patchy particle enzymes, with the hard sphere enzyme surface represented in blue while the active site patches are represented by smaller spheres (red, yellow and purple circles) on the surface of the hard sphere. The central hard spheres (blue circle in Fig.1A) representing the enzyme surface are larger in diameter than the active sites (red, yellow and purple circles in Fig.1A). The enzyme surface (blue color hard spheres in Fig.1A) are involved in isotropic interactions with other proteins as well as non-specific interactions with ligands at interaction strengths of *ε*_*protein*–*protein*_ and *ε*_*protein*–*ligand*_, respectively. While non-specific interactions between an enzyme and a ligand may occur anywhere on the surface of an enzyme, a reaction can occur only when an active site comes in contact with its complementary ligand.

**Figure 1.**
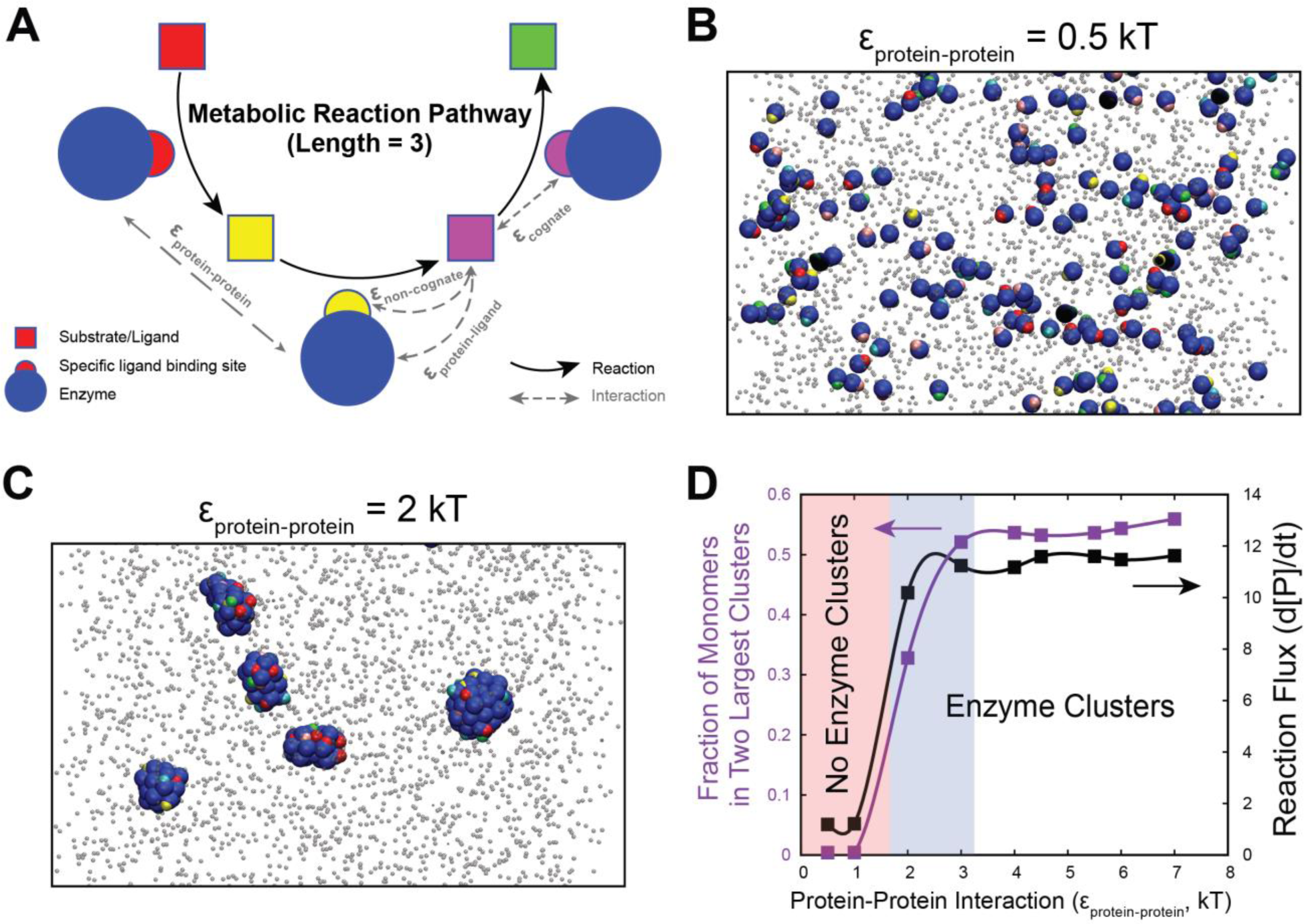
Clustering of enzymes. (A) Schematic figure of the 3-reaction pathway simulated using LD simulations of patchy particle enzymes. A reaction can occur with a probability (*P*_*react*_) when the complementary substrate binds to the active site. The active site is modeled as a patch on the surface of the enzyme such that only a small fraction of the enzyme surface belongs to the active site. Enzymes can interact with other enzymes with an attractive potential of strength ε_protein-protein._ Ligands interact with the surface of the protein (ε_protein-ligand_) or with non-cognate active sites with a strength ε_non-cognate._ When the active site is bound to the complementary, cognate ligand, the strength of the interaction is ε_cognate._ (B) and (C) represent the state of the multi-particle 3-enzyme system for protein-protein interaction strength of 0.5 kT and 2 kT, respectively. The enzymes are shown as large blue-colored spheres with side patches of different colors (representing different enzymes along the 3-reaction pathway). The substrate molecules are shown as smaller grey spheres. At protein-protein interaction strength of 0.5 kT, the enzymes persist in monomeric state and the reaction occurs in the bulk. On the other hand, at a protein- protein interaction strength of 2 kT, the enzymes exist in the form of multi-enzyme clusters and the reactions occur within these enzyme clusters. (D) Cluster sizes and corresponding reaction flux as a function of protein-protein interaction strength (ε_protein-protein_) in LD simulations. The most significant increase in reaction flux is observed at interaction strengths that favor large clusters.

Here we consider a linear enzymatic reaction pathway composed of *l* chain reactions as shown by Eq. 1 (also see Fig. 1A). Catalyzed by enzyme *E*_*i*_, substrate *S*_*i*−1_ can be converted to *S*_*i*_ inside the cluster. After *l* chain reactions, substrate *S*_0_ will be converted to the final product *P*.

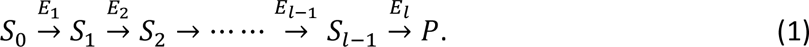

In order to model this enzymatic pathways, we employ a stochastic reaction-diffusion modeling protocol as follows. Whenever a substrate/intermediate is bound to the cognate, active site specific to it, a reaction can occur with probability *P*_*react*_. In our simulations, we vary this probability to model enzymes with different catalytic efficiencies. By modulating the enzymatic reaction rates, which is governed by reaction probability *P*_*react*_, we can simulate enzymatic activities ranging from the reaction-limited to the diffusion-limited regimes.

The active site patches (red, yellow and purple semicircles in Fig.1A) can engage in two distinct modes of interaction: cognate and non-cognate binding. Cognate binding (interaction strength of *ε*_*cognate*_) denotes the interaction between a cognate active site of an enzyme and its complementary ligand, as each ligand-binding site at the active site of an enzyme is specific to one particular substrate (or intermediate) along a linear pathway (see Eqn. 1). Non-cognate binding at an interaction strength of *ε*_*non*–*cognate*_ is defined as interactions between an active site of an enzyme and other non-cognate substrates. In our simulations, *ε*_*non*−*cognate*_ is equal to *ε*_*protein*−*ligand*_ meaning that interactions between non-complementary ligand and active site are treated the same way as non-specific interactions between the protein surface and ligand molecules.

#### Potential Functions and Simulation Variables

Interactions between any pair of enzymes (i.e., *ε*_*protein*−*protein*_), protein-ligand interactions outside of the binding site (i.e., *ε*_*protein*−*ligand*_), as well as non-cognate binding (*ε*_*non*−*cognate*_) are all modeled using the Lennard-Jones (LJ) potential,

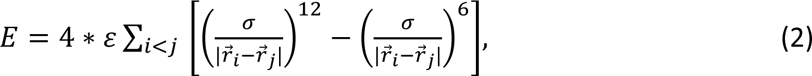

for all 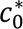. Here, *σ* is the diameter of central body particles (σ_enz_ for enzyme surface), which is set to 20 Å. The diameter of the patch (σ_patch_) used to model the active site of the protein was set to 5 A, same as that of the ligand molecules σ_ligand_. *r*_*c*_ refers to the interaction cutoff beyond which LJ potential are neglected. The cutoff *r*_*c*_ is set to 2.5 times of *σ*, for the LJ potential. The strength of the attractive part of the LJ potential, *ε*, is tuned to vary the strength of different interactions as shown in results part. As reported above, depending on the identity of the interacting particles, *ε* takes one of the following values -- *ε*_*protein*−*protein*_ (between proteins) or *ε*_*protein*−*ligand*_ (between ligands and the protein surface outside of active site).

Cognate binding (*E*_*cognate*_) is defined by a continuous square-well potential,

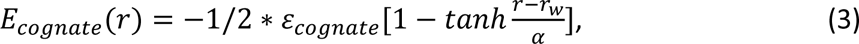

for all *r* < *r*_*w*_, where *r*_*w*_ is the interaction cutoff beyond which cognate binding are neglected. The strength of cognate binding between the correct, complementary substrate and the binding site *ε*_*cognate*_ is set to 4 kT whereas the interaction cutoff *r*_*w*_ is set to 0.12*σ*, where *σ* is the size of the central hard sphere that the binding site is part of. This *r*_*w*_ of 0.12*σ* ensures a valency of one per attractive site. The range of the attractive interactions between the active site and the complementary ligand, is, therefore, roughly equal to 2.5 A, mimicking typical hydrogen bond lengthscales in biomolecular systems.

We used the LAMMPS molecular dynamics package ^15^ to perform the dynamic simulations, where the simulator solves Newton’s equations with viscous force, and a Langevin thermostat ensuring an NVT ensemble with temperature *T* = 310 K. An integration timestep (*dt*) of 15 fs was used for the simulations. The viscosity of the medium was set to that of water. As these patchy particle proteins are defined as a multi-centre rigid body with no internal dynamics in our simulations, we use the rigid particle definition in LAMMPS ^15^.

### Semi-analytical Reaction-Diffusion Model

In order to probe a wider range of enzymatic rates as well as enzymatic reaction pathway lengths, we employ a variation of the semi-analytical reaction-diffusion model first proposed by Castellana et al ^6^. A representative enzymatic reaction pathway with *l* sequential reactions is shown in Eq. 3. In this model, we assume that cell can be divided into several identical spherical regions (referred to as “reaction unit” thereafter, outlined by cyan circle in Fig. 2C), which is composed of one enzyme cluster positioned in the center (green circle in Fig. 2C) and the surroundings (white region between cyan and green circle). All the enzymes (*E*_*i*_) are concentrated inside the cluster while the substrates (*S*_*i*_) can diffuse between cluster and its surroundings. Corresponding reaction-diffusion equations can be written as follows.

**Figure 2.**
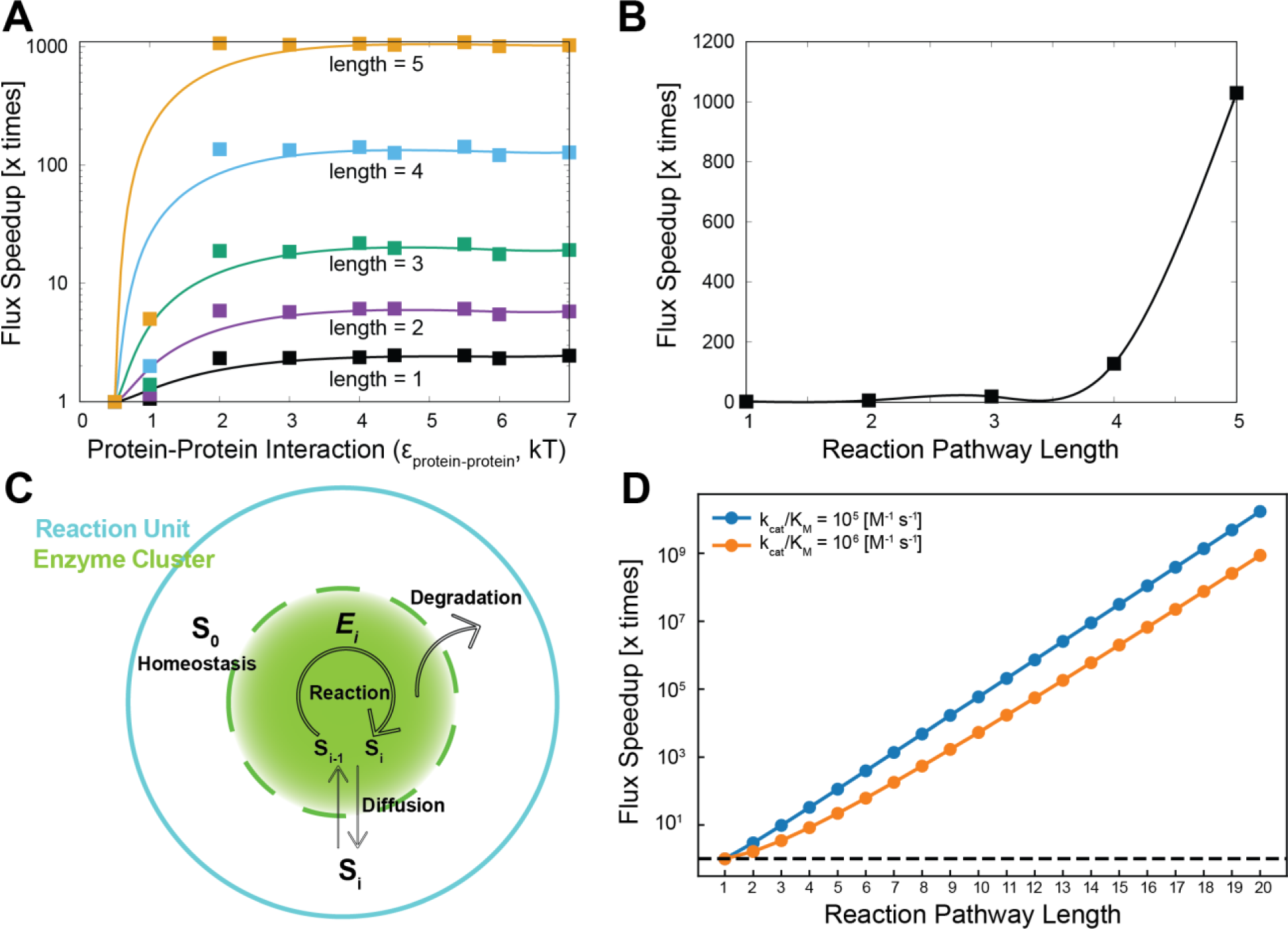
The effect of pathway lengths. A) Speedup in reaction flux (clustered vs. delocalized) in LD simulations at different pathway lengths for different strengths of protein-protein interaction. Each curve represents a pathway length. As the length of the pathway grows, we see a significant speedup in flux over the bulk reaction scenario. The solid curves are a guide to the eye. B) Speedup in reaction flux in LD simulations as a function of pathway lengths, for protein-protein interaction strength of 2 kT. (C) Schematic figure of reaction-diffusion model. Cell cytoplasm can be divided into multiple reaction units (outlined by cyan circle), each of which contains one enzyme cluster (green) and the surroundings (white). Enzymes are all concentrated in center cluster, where metabolic reactions happen. Substrates can diffuse between cluster and the surroundings within reaction unit (see methods). Substrates that have not reacted within the degradation timescale (see degradation rate β in methods) are degraded. The concentration of initial substrate *S*_0_ is maintained through homeostasis by upstream reactions. (D) Reaction- diffusion model prediction of speedup in reaction flux (clustered vs. delocalized) in a broad range of metabolic reaction pathway lengths. Here we use the reaction-limited enzyme with *k*_*cat*_/*K*_*M*_ of 10^5^ and 10^6^ M^−1^ s^−1^ as examples.

The fate of the first substrate in the pathway, S_0_, the diffusion-reaction equation can be described as,

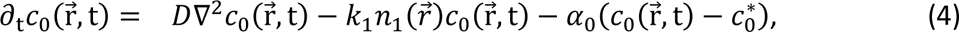

where the first term represents the flux of the initial substrate *S*_0_ into the cluster while the second term describes the conversion of S_0_ to the first intermediate S_1_. The last term represents concentration of initial substrate *S*_0_ which is controlled to a homeostatic concentration *c*^∗^ via upstream metabolism reactions which occur at rate *α*_0_.

The fate of all other intermediates along the pathway can be written by the following set of reactions described in Eq. 5,

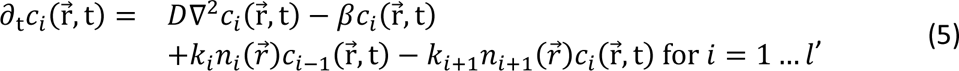

where *β* in the above equation represents the rate at which intermediates (*S*_1_ to *S*_*l*_) get degraded if they do not get converted to the downstream substrate before the characteristic degradation timescale defined by the degradation rate *β*. In Eq. 4 and 5, *D* is the diffusion coefficient while *k*_*i*_ is enzyme’s catalytic efficiency (i.e., *k*_*cat*_/*K*_*M*_, *k*_*cat*_ is the catalytic rate constant, *K*_*M*_ is Michaelis constant). 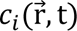 represents the concentration of substrate *S*_*i*_ at position 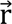 and time *t* while 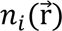 is concentration of enzyme *E*_*i*_ at position 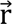. For the sake of analytical tractability, we assume that the diffusion coefficient for all substrates are equal.

In order to ascertain the net gain or decrease in reaction flux upon enzyme clustering, we also study the delocalized scenario, in which enzymes are distributed uniformly across a larger reaction cell (low local density) -- representing the cytoplasm -- rather than condensing into a single cluster (high local density of enzymes). All the enzymes (*E*_*i*_) and substrates (*S*_*i*_) can diffuse within the whole reaction unit. The enzymatic reaction pathway, degradation and homeostasis are the same as in the clustered scenario mentioned above.

#### Assumptions of the semi-analytical model

We first assume that reaction cell – representing whole cytoplasm - can be divided into several identical spherical regions. Each of those spherical regions is defined as a reaction unit (outlined by cyan circle in Fig. 2C), which contains one enzyme cluster (green circle in Fig. 2C) and its surrounding region (white region between cyan and green circle). It was also assumed that in the region surrounding the reaction unit, there is no enzyme. The substrates, however, can diffuse between the cluster and it surroundings. The density of enzymes within a multi-enzyme cluster is assumed to be uniform. In the delocalized scenario, where the enzymes exist as monomers, we assume that the same number of enzymes as in the cluster are uniformly distributed across the larger volume of the entire reaction unit. These assumptions were previously used by Castellana et al^6^ to study the influence on enzymatic clustering on multi-reaction pathways. We also assume spherical symmetry wherein the concentration of substrates *c*_*i*_ depends only on radius *r* – the radial distance from the center of cluster. For analytical simplification, the catalytic efficiency for individual enzymes, *k*_*i*_ was considered to be the same for all steps in the pathway, and will be hereby referred to as *k* for all reactions. The density of enzyme *i* is defined as *n*_*i*_ = *n*_*all*_/*l* (referred to as *n* for all enzymes thereafter), where *n*_*all*_ is the total enzyme density within the reaction unit. In other words, all enzymes in the pathway have the same density and operate at the same catalytic efficiency.

#### Boundary conditions and analytical solutions

To solve the reaction-diffusion model numerically, we adopt a no-flux boundary condition where substrates cannot diffuse out of the reaction unit.

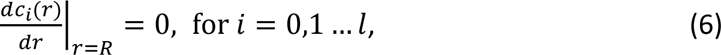

where *R* is the radius of the reaction unit. The total density of enzymes 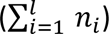 within a cluster cannot exceed the maximum permissible density (*n*_*max*_) in order to account for excluded volume interactions.

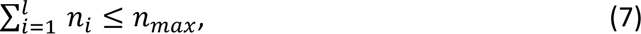

The pathway efficiency, *η*, is defined as the fraction of substrate population *S*_0_ that is being converted to product when all the reactions reach the steady state (i.e., left hand of Eq. 4 and 5 equal to 0). Note that the pathway efficiency ‘*η*′ here is not to be confused with enzyme catalytic efficiency ‘k’ which corresponds to the catalytic efficiency of individual enzymes.

The pathway efficiency *η* for enzymatic reaction pathway with *l* chain reactions can be defined as follows,

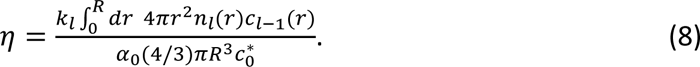

The numerator in Eq. 8 defines the rate of production *P* formation. The denominator describes the maximal possible supply rate of substrate *S*_0_ by homeostasis when there is no substrate *S*_0_ in reaction unit. Therefore, *η* is a time-independent steady-state quantity.

For mathematical tractability, we assume that substrates are also uniformly distributed inside both the cluster and the surroundings, through which we are able to obtain the analytical solution for pathway efficiency *η* and qualitatively illustrate the benefits of enzyme clustering for reaction pathway with different enzyme catalytic efficiencies *k*. For the delocalized case where the enzymes exist in monomeric form in the reaction cell, pathway efficiency *η*_delocalized_ can be analytically solved as follows (See Supplementary text for derivation),

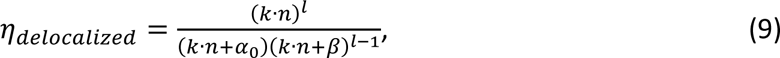

where *k* is the enzyme catalytic efficiency for all reactions in the pathway, *n* is the density of any enzyme in the pathway based on the maximal density set by the relationship in Eq. 7.

Weak non-specific interactions between enzymes and substrates, *ε*_*protein*−*ligand*_, are accounted via the following boundary conditions which define the concentration of substrates in the enzyme cluster and the surroundings,

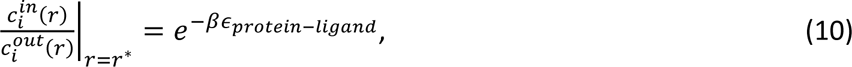

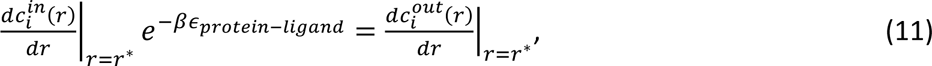

where *in* and *out* denote enzyme cluster and the surroundings respectively, *r*^∗^ is the radius of enzyme cluster, *i* ranges from 2 to *l* as the concentration of initial substrate *S*_0_ is controlled to a homeostatic concentration 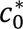 at rate *α*_0_ through its upstream metabolism reactions for both inside and outside enzyme cluster. In enzyme cluster, efficiency of the first reaction in the pathway *η*_1_ is given as ^6^

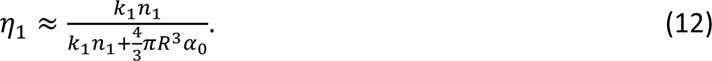

For all other reactions in the pathway, (2 ≤ *i* ≤ *l*),

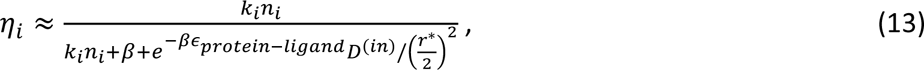

where *e*^−*βεns*^ is added based on ref. 6 so to account for weak non-specific interactions *ε*_*protein*−*ligand*_ between enzymes and substrates. It must be noted that this weak non-specific interaction allows substrates to dwell longer in the cluster, thereby decreasing the flux of substrates out of the cluster. This phenomenon was systematically studied using LD simulations, the results of which are discussed later in this article.

The pathway efficiency for the whole reaction pathway can be obtained through

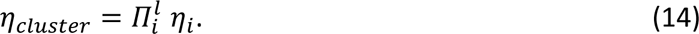

Finally, the metabolic flux speedup, the ratio between reaction flux of enzyme cluster and that of delocalized scenario, can be formulated as follows,

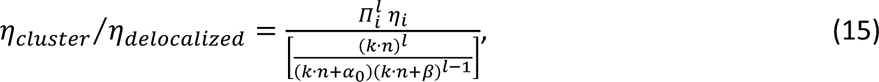

where *η*_*i*_ is given by Eq. 12 and Eq. 13.

All the numerical solutions are based on experimental data for purinosome ^4, 10^. The size of reaction unit *R* for purinosome is 1.3 *μm*. The size of cluster *r*^∗^ for purinosome is 0.3 *μm*. Diffusion coefficient in enzyme clusters is 1 *μm*^2^*s*^−1^. Diffusion coefficient outside enzyme clusters or in delocalized case is 10 *μm*^2^*s*^−1^. Degradation rate for all substrates is 1 *s*^−1^ ^6^. *n*_*max*_ is calculated as 25 *mM*, which assumes single enzyme size as 2 *nm* and enzymes take 50% volume fraction at most due to the water solvation shell on the surface of enzymes ^6^. The total enzyme density *n*_*all*_ within the reaction unit is fixed as 1000 *μm*^−3^, which satisfies the Eq. 7. The density of enzyme *i* is defined as *n* = *n*_*all*_/*l*. *α*_0_ is set as 0.1 *s*^−1^ ^6^. *ε*_*protein*−*ligand*_ is set as 0.7 kT to match the simution.

### Bioinformatics Study

We first obtain the curated database of enzyme catalytic efficiencies from Ref. 11, which exclusively includes natural enzymes and reactions of natural substrates. Metabolic pathways associated with each enzyme are identified through their reaction IDs via KEGG database ^21^ (https://www.kegg.jp/kegg/). Given the multiplicity of pathways in which one enzyme typically participates, the average enzyme counts across all associated pathways are computed and used to represent the size of enzyme clusters that they belong to. Database and detailed workflow (Fig. S6) are provided as Supplementary Materials.

#### Statistical analysis

The statistical significance of the differences between the two groups (i.e., p-value) was evaluated using an independent two-sample t-test. The t-test is based on the student’s t-distribution, and its null hypothesis posits no difference in means between the two groups. A p-value is computed under this null hypothesis, with a smaller p-value indicating stronger evidence against the null.

## Results

### Clustering of enzymes in long reaction pathways can improve reaction efficiency over bulk by several orders of magnitude

Clustering of enzymes and the accompanying density transitions have been experimentally reported to improve intermediate processing and the rate of product formation ^6^. Here, we first use diffusion-reaction Langevin Dynamics (LD) simulations (see Methods section for details) to study the effect of protein clustering on the rate of product formation for a linear three-reaction enzyme pathway (Fig. 1A). In this three-enzyme system, we begin with the initial substrates (red square in Fig. 1A). When the active site (red semicircle in Fig. 1A) of the first enzyme comes in contact with the initial substrate, a reaction occurs with a probability *P*_*react*_ and the first intermediate (yellow squares in Fig. 1A) in the pathway is produced. This scheme can be generalized for all subsequent reactions till the product P (green square in Fig. 1A) is formed. In order to study the effect of enzyme clustering on the efficiency of the three-reaction pathway, we systematically vary the strength of protein-protein interactions (ε_protein-protein_), keeping all other parameters the same. In Fig.1B and 1C, we show the state of the system for two different strengths of ε_protein-protein_ – 0.5 kT and 2 kT—respectively. As seen from Fig. 1B and 1D, the enzymes persist in a monomeric state at a weak interaction strength of 0.5 kT, with the reactions therefore occuring in the bulk. An increase in interaction strength from 0.5 to 2 kT results in formation of multi-enzyme clusters, with the reactions now occuring within the enzyme clusters. It must, however, be noted that the critical value of interaction strength would depend on the concentration of enzymes in the system. In the results reported in the current work, all simulations were performed at a volume fraction of 0.05, i.e 5% of total volume in the simulation was occupied. This number is within the biologically relevant range of occupied volume fraction in the cell ^16, 17^.

In Fig. 1D, we plot the extent of clustering and reaction flux (d[P]/dt, the change of product concentration with time) as a function of protein-protein interaction strengths. Note that the flux plotted here is for the final product of the 3-reaction pathway. In order to compute the extent of clustering, we use the number of enzyme molecules in the two largest multi-enzyme clusters in the simulation box. As evident from Fig. 1D, below an interaction strength of 2 kT, the enzymes exist in monomeric state and all reactions fluxes are low. For interaction strengths above 2 kT, the clustered state of enzymes is favored since we observe several orders of magnitude increase in reaction flux in the LD simulations (Fig.1C). The extent of clustering and reaction flux curves in Fig. 1D suggest that speedup in reaction flux is indeed a function of clustering of enzymes at interaction strengths of 2 kT and higher. Importantly we observe the speedup in metabolc flux to occur concommitantly with phase transition leading to spatial clustering of enzymes.

### Longer pathways gain siginficantly from clustering

So far, we employed LD simuations with physical descriptions of tunable, protein-protein interactions to study the effect of enzyme clustering on reaction flux in a three-reaction pathway. However, biochemical pathways could be several reactions in length. For instance, the purine biosynthesis is a ten-reaction pathway ^4, 20^. Therefore, we further probe whether the gain in reaction flux upon enzyme clustering persists for longer pathways. We first extend the reaction scheme in our LD simulation from 3 (Fig. 1A) to 5 reactions and systematically study the reaction flux for different pathway lengths. In Fig. 2A, we plot the speedup in reaction flux as a function of protein-protein interaction strength, for pathways of different lengths. As evident from Fig. 2A and B, as the pathway length becomes longer, the speedup in reaction flux upon clustering increases dramatically compared to the bulk reaction scenario. As we approach pathway lengths of 5, we observe 3-orders of magnitude gain in reaction flux (d[P]/dt) over the bulk (Fig. 2A and 2B). In Supplementary Fig. S1 (panels A-E), we also plot the flux vs. substrate concentration profiles for the 5 reactions along the linear pathway, at different strengths of protein-protein interactions to modulate the extent of enzymatic clustering. As seen Fig. S1A-E, an increase in protein-protein interactions from 1-2 kT results in an increase in reaction flux, for all the 5 reactions along the linear pathway. However, for protein-protein interactions stronger than 2kT, the relative speedup in reaction flux (*ε*_*protein*−*protein*_ ≥ 2kT vs. < 2 kT) is progressively higher for enzymes further along the reaction pathway. Interestingly, while the flux vs. substrate concentration profiles for the first and second reactions (Fig. S1A and S1B) are monotonically linear in the range of substrate concentrations simulated, reactions 3 to 5 (Fig. S1C to E) display signatures of saturation at higher substrate concentrations. Clustering of enzymes allows the system to reach saturation fluxes for the latter reactions in the pathway, without actually altering their microscopic properties.

While the LD simulations suggest that the speedup in reaction flux is higher for longer pathways, these simulations cannot access long pathway lengths. To that end, we employ a semi-analytical reaction-diffusion model adapted from Castellana et al. ^6^ (Fig. 2C, also see Methods section) to systematically study reaction flux speedup for broader range of metabolic pathway lengths. In Fig. 2D, we plot the metabolic reaction flux speedup (Eq. 15) due to clustering, for varying pathway lengths. The reaction flux speedup is computed as follows, for varying lengths *l* of the pathway

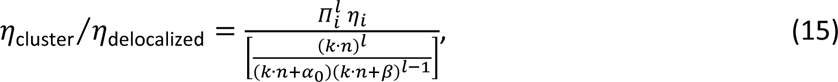

where *η*_*i*_ is given by Eq. 12 and Eq. 13 (see Methods). Consistent with the LD simulations, the results from the analytical model also show that the flux speedup ratio increases with increasing pathway length. Therefore, our results indicate that longer reaction chains gain the most from enzymatic clustering (Fig. 2), suggesting that it could be a plausible mechanism to improve efficiency of long biochemical pathways.

### Weak non-specific protein-ligand interactions are crucial for flux speedup in enzyme clusters

Active site residues make up only a fraction of the enzyme surface. A mere collision of the substrate with the enzyme would therefore not be a sufficient condition if the substrate is unable to find the binding site before diffusing away from the enzyme. In this context, diffusion of ligands on the protein surface (after the initial collision) has been shown to enhance the process of ligand’s search for the binding site^18^. Given that the binding site of enzymes in our simulation comprises only a small fraction of the surface, the mean dwelling times for ligands in the vicinity of the protein could dictate the fate of a protein-ligand encounter.

We therefore systematically vary the strength of non-specific protein-ligand interactions (ε_protein-ligand_), across all substrate-enzyme pairs and across the hard-sphere enzyme surface. As we increase ε_protein-ligand_ from 0 to 2 kT, we observe about two orders of magnitude increase in reaction flux speedup, at fixed protein-protein interaction strength of 2 kT above the threshold of enzyme clustering (Fig. 3A, purple curve). Below a critical non-specific protein-ligand interaction strength, we observe negligible product formation at simulation timescales (Fig. 3A, purple curve). Interestingly, as we increase the strength of non-specific protein-ligand interactions above 2 kT, we see a dropoff in reaction fluxes, suggesting that there exists an optimal range of non-specific protein-ligand interactions that results in the highest product formation rates. In the regime of weak non-specific protein-ligand interactions the ligands exhibit free-diffusion in solution. In the regime where non-specific interactions result in the highest speedup in reaction flux (green-shaded region in Fig. 3), we see a 5-fold decrease in diffusion coefficient of the ligand suggesting that the ligands dwell in the enzyme cluster while continuing to remain diffusive. On the other hand, as non-specific interaction strength increases to 3 kT and beyond (red-shaded region in Fig. 3), there is a dramatic loss of diffusivity of the ligand, suggesting that the ligands get sequestered onto the enzyme surfaces in this regime. To probe the origins of this dropoff in efficiency, we also calculate the fraction of ligand molecules in an enzyme-bound state (Fig. S2). As we increase the strength of non-specific protein-ligand interaction from 0 to 2 kT, the ligands populate the enzyme clusters at higher fractions (Fig. S2, black curve). However, further increase in interaction strength results in almost all substrates being bound to the enzyme surfaces (Fig. S2 black curve). In this regime (ε_non-cognate_ > 2 kT), diffusivity of the enzyme-surface bound substrates drops two-orders of magnitude sower within the enzyme clusters, slowing down the process of finding the correct active site. We also determined the fraction of ligands bound to the correct binding site as a function of non-specific protein-ligand interaction strength (Fig.3B) to check whether occlusion of binding sites is another reason for the dropoff in reaction efficiency. Interestingly, while we observe negligible active site binding by non-cognate substrates in the weak ε_protein-ligand_ regime (white region in Fig. 3B), in the regime of optimal ε_protein-ligand_ strength, the fraction active sites bound to their cognate substrates far exceeds that of active sites with bound non-cognate substrates. As the strength of non-specific protein-substrate interactions exceeds 3kT, a point where specific (for cognate binding) and non-specific protein-ligand interactions are of equal strength, the active sites get significantly occluded by incorrectly bound substrates which do not result in a reaction. A combination of the dropoff in diffusivity of substrate within the cluster as well as occlusion of binding sites at high non-specific protein-ligand interactions results in the non-monotonic dependence of reaction flux on non-specific ligand protein interactions. Crucially, in the absence of non-specific protein-ligand interactions, there is no speedup in reaction flux upon enzyme clustering. Nerukh et al. ^18^ report a non-specific surface interaction of 1.6-2 kT between the DHFR surface and TMP that results in optimal ligand-binding kinetics, consistent with our findings.

**Figure 3.**
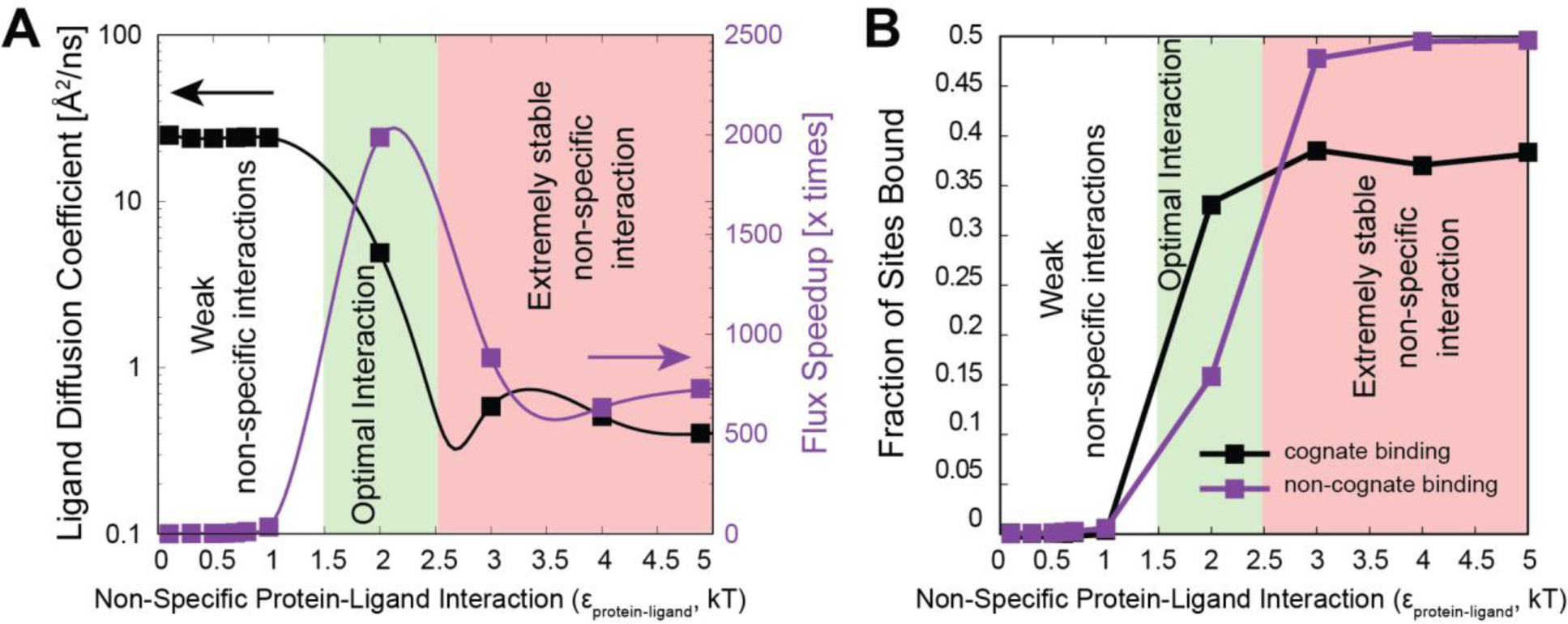
Importance of non-specific protein-ligand interactions. A) In the purple curve, speedup in reaction flux shows a dramatic increase upon introduction of non-specific protein-ligand interactions. A further increase in non-specific protein-ligand interactions results in a dropoff in flux. In the black curve, we plot the diffusion coefficients of the ligand as a function of *ε*_*protein*−*ligand*_. In the regime where we see a dropoff in reaction flux, the ligand molecules show an order of magnitude decrease in diffusion coefficient, suggesting a loss of diffusivity in this regime. B) Fraction of sites bound by cognate or non-cognate ligands at different non-specific protein-ligand interactions. When *ε*_*protein*−*ligand*_ = 2 kT, binding sites are mainly occupied by cognate ligands while beyond 2 kT, non-cognate binding dominates and very few sites are left available.

To further understand how non-specific protein-ligand interactions affect substrate dwell times on the enzyme, we simulate a 2 particle, enzyme-substrate system wherein the enzyme and substrate are freely diffusing particles that encounter each other in solution to undergo a reaction (Fig. S3). Using LD simulations of patchy particle enzymes and substrates as outlined previously (Fig. 1A and Methods), we vary the strength of non-specific protein-ligand interactions and compute the key processes that govern a reaction – i) The mean dwell time, the time spent by the ligand in contact with the enzyme before it diffuses away to solution (t_dwell_), ii) the mean first passage time for the ligand to find the active site on the enzyme upon coming in contact with the enzyme surface (t_find_), and iii) the mean first passage time to go from the free state to formation of the product (t_react_). In Supplementary Fig. S3A, we plot the dwell time as a function of non-specific protein-ligand interaction strength. As evident from Fig. S3A (black curve), non-specific protein-ligand interaction result in an increase in dwell times allowing the ligand to stay in the vicinity of the protein longer before diffusing away. Crucially, the increase in dwell times is accompanied by a concomitant decrease in the time taken for the ligand to find the binding site (t_find_) on the enzyme (Fig.S3B, purple curve). This speedup in t_find_ results in an overall reduction in the mean first passage time to go from the unbound state to product formation (t_react_ in Fig. S3B). It must however be noted that unlike the multi-enzyme, long reaction pathway scheme, the increase in non-specific protein-ligand interaction in a one-reaction enzyme-substrate scheme does not result in a dropoff in reaction flux. This could be attributed to the fact that in the presence of strong non-specific interactions in a multi-reaction, multi-enzyme pathway could result in occlusion of binding sites by the non-cognate substrate resulting in slower reaction flux. The two particle system helps us effectively demonstrate how reaction fluxes benefit from an increase in dwelling time facilitated by non-specific protein-ligand interactions. However, the dropoff in turnover at higher ε_protein-ligand_ is a consequence of active site occlusion by non-cognate ligands, a signature of multi-reaction, multi-enzyme clusters.

Therefore, weak non-specific interactions between ligand and protein surfaces can allow the ligands to dwell long enough in the vicinity of the protein, allowing them to effectively reach the ligand-binding site where reaction occurs, resulting in more productive collisions. In the absence of such a weak attractive forcfield in the vicinity of the enzyme cluster, the ligands would be unable to “search” the cluster effectively for the cognate active sites where reactions occur before diffusing away from the cluster.

In our simulations so far, we consider all protein-protein interactions to be homogenous and isotropic. In other words, there is a uniform spatial distribution of enzymes within the cluster. However, biological interactions could display heterogeneity. Further, tunneling of substrate from one enzyme to another could also result in a gain in specificity and efficiency of enzyme clusters. In this context, we look at a scenario where successive enzymes in the pathway interact at different interaction strengths *ε*_*neigh*_ while all other pairwise protein-protein interactions *ε*_*non*−*neigh*_ are set to 2 kT, a regime where clusters are favored (Fig. 1C). Unlike the gain in reaction flux upon formation of enzyme clusters (Fig. 1C, at *ε*_*protein*−*protein*_ ≥ 2 kT) and the introduction of weak non-specific protein-ligand interactions (Fig. 3), there is no significant gain (Fig. S4A-E) evident from introducing stronger favorable interactions between successive enzymes of the pathway. Therefore, weak homogenous non-specific interactions can sufficiently accelerate long reaction pathways even in the absence of specific fine tuned protein-protein interaction networks within metabolons.

### Clustering of enzymes provides dramatic gains for “inefficient” reaction-limited enzymes while providing neglibile gains for diffusion-limited enzymes

In our results so far, we probed how protein-protein and protein-ligand interactions could influence the rate of product formation, for varying pathway lengths. However, enzymes could also vary significantly in their catalytic efficiencies, with the “average” enzyme being several orders of magnitude slower than the “ideal” diffusion-limited enzyme^11^. To probe whether clustering influences reation-limited and diffusion-limited enzymes differentially, we first vary the probability of reaction in order to tune catalytic efficiency in Langevin dynamics simulations. In Fig.4A, we plot the speedup in reaction flux (d[P]/dt) compared to the unclustered monomeric state in the bulk. For diffusion-limited enzymes, we observe no speedup over bulk reaction fluxes, across the range of protein-protein interaction strengths studied (Fig. 4A, black curve). However, for enzymes with less than optimal catalytic efficiencies, where reaction is the rate-limiting step (purple and green curves in Fig. 4A), we observe a dramatic speedup in reaction fluxes at protein-protein interaction strength of 2 kT or higher corresponding to the onset of clustering. The magnitude of this speedup also shows a dependence on the presence or absence of weak non-specific protein-ligand interactions (Fig. 4A green vs purple curve. also see Fig. 3). For protein-protein interaction strengths above 2 kT where the clustered state of enzymes is favored (Fig. 1A), we observe 2 distinct regimes of clustering-associated speedup in reaction fluxes (Fig. 4B). In the regime where the reaction timescales are much slower than the diffusion timescale (*k*_*react*_ ≪ *k*_*diff*_ region in Fig.4B), we observe the final reaction in the 3-reaction pathway speedup by an order-two orders of magnitude over that of the bulk. On the other hand, when the reaction timescale is of the same order as that of diffusion (*k*_*react*_ = *k*_*diff*_ region in Fig.4B), there is no speedup over the bulk reaction fluxes. At intermediate reaction rates (*k*_*react*_ < *k*_*diff*_ region in Fig.4B), we observe a marginal speedup in product formation.

**Figure 4.**
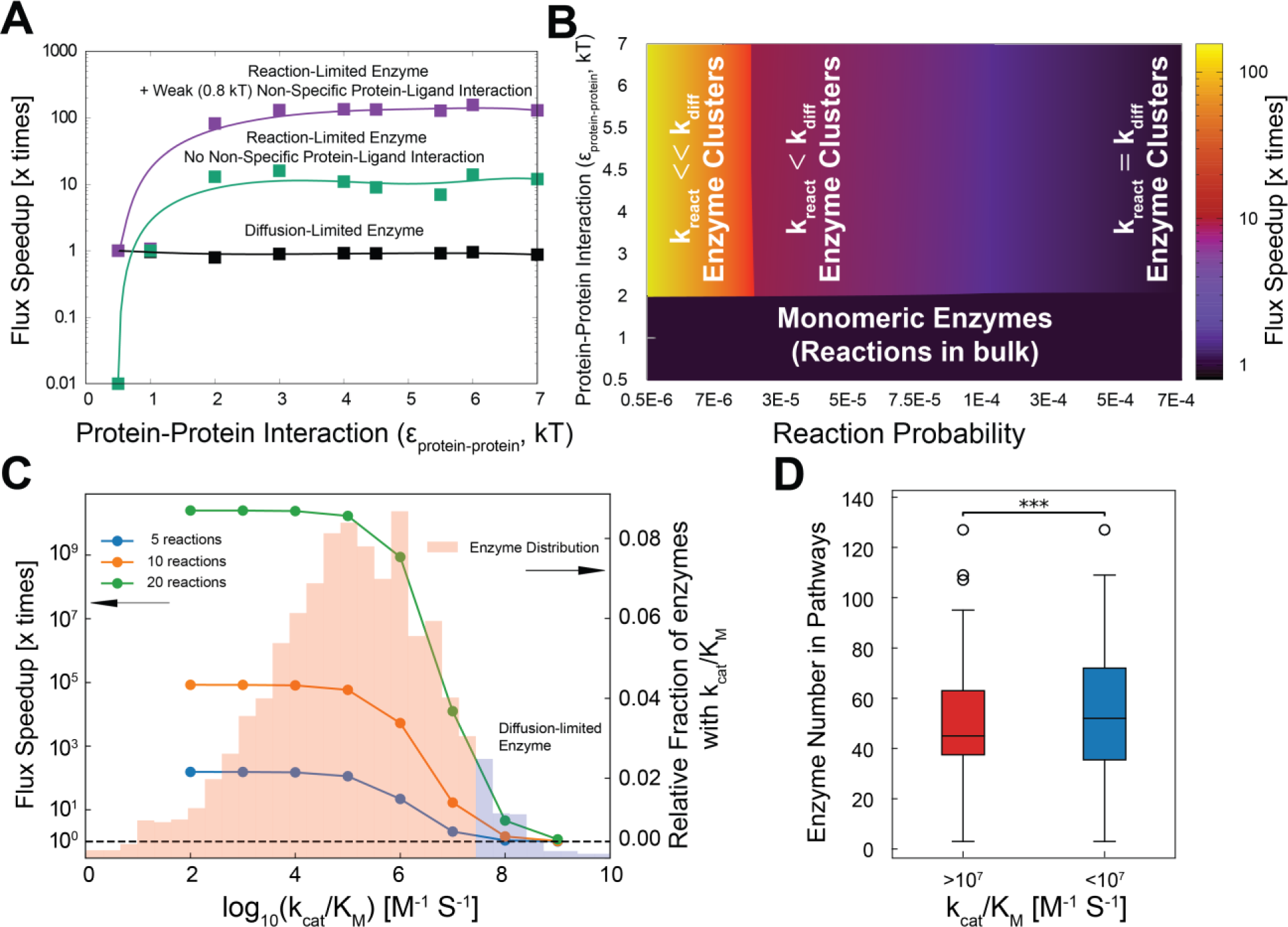
Reaction speedup for diffusion and reaction-limited enzymes. A) Speedup of reaction flux in enzyme clusters compared to reactions occuring in bulk, at same enzyme and substrate concentrations. Reaction-limited enzymes (green and purple curves) show a dramatic increase in speedup at protein-protein interactions of 2 kT or higher corresponding to the phase transition leading to the onset of clustering. Diffusion-limited enzymes (black curve), on the other hand, show no such speedup across protein-protein interaction strengths studied. Note that the results plotted here are for a pathway length of 3 reactions. The reaction fluxes also gain from the presence of a weak non-specific interaction (purple curve). B) Phase diagram showing the speedup over bulk reaction fluxes, for varying protein-protein interaction strengths and reaction probabilities. The most significant speedup is observed for reaction-limited enzymes. Color code from black to yellow denotes the increase of reaction flux speedup. *k*_*react*_ and *k*_*diff*_ represent reaction rate and diffusion rate with unit 1⁄*s*, respectively. C) Reaction flux enhancement due to enzyme clustering compared with delocalized enzyme system (i.e., enzymes distributed uniformly inside the surroundings) for different reaction pathway lengths at different catalytic efficiencies *k* (i.e., *k*_*cat*_/*K*_*M*_). D) Average enzyme number in pathways for different catalytic efficiencies *k* (i.e., *k*_*cat*_/*K*_*M*_). 3-star (***) means p-value is less than 0.001. The boxplot displays the interquartile range (IQR) with median marked. Whiskers extend 1.5 times of the IQR, and points beyond are outliers. In agreement with the theory, less efficient enzymes populate longer pathways than highly efficient diffusion-limited enzymes.

We further systematically vary catalytic efficiencies (*k*_*cat*_/*K*_*M*_) in the semi-analytical model to probe the whole range of *k*_*cat*_/*K*_*M*_ observed in naturally occuring enzymes (see distribution of *k*_*cat*_/*K*_*M*_ in the background in Fig. 4C). As highlighted in the seminal study by Bar-Even et al ^11^, the distribution of *k*_*cat*_/*K*_*M*_ in prokaryotic and eukaryotic enzymes peaks at 4-5 orders of magnitude lower than the diffusion-limit (Fig. 4C, red-shaded background). Crucially, from the semi-analytical calculations, the biggest gain in reaction efficiency upon clustering was observed for catalytic efficiencies which correspond to the reaction-limited regime of the distribution in Fig. 4C. In the regime of *k*_*cat*_/*K*_*M*_ corresponding to the right-handed tail of the distribution, we see a dramatic dropoff in efficiency gain upon clustering. This suggests that the clustering of enzymes significantly increases the frequency of collisions between substrates and enzymes before substrates diffuse away or get degraded resulting in an enhancement in reaction efficiency. The acceleration of reaction flux for reaction-limited enzymes mainly comes from drastic decrease in efficiency in delocalized systems as enzymatic pathways become longer. For long pathways, in the delocalized case, the mean free path for the intermediate to find the next enzyme in the pathway is much longer compared to the clustered state. For diffusion-limited enzymes, fewer collisions are required for a productive encounter, as compared to reaction-limited enzymes. The rate-determining step, is therefore, the diffusion process leading to first encounter between active site of an enzyme enzyme and substrate and therefore we observe no significant gain in efficiency upon clustering in this regime. Since reactions proceed fast in diffusion-limited enzymes, no sharp decline of efficiency is shown in delocalized systems. Therefore, the efficiency enhancement ratio is close to 1 in the extreme diffusion-limited case (*k*_*cat*_/*K*_*M*_ = 10^9^*M*^−1^*s*^−1^). Overall, these results suggest that clustering of enzymes could dramatically increase the reaction flux of reaction-limited enzymes by increasing the local density of enzymes.

Our LD simulations and semi-analytical calculations suggest that enzymes with weaker catalytic efficiencies belonging to longer pathways benefit the most from enzymatic clustering. On the other hand, enzymes with higher catalytic efficiencies do not benefit from clustering and therefore should tend to work in isolation or belong to shorter pathways. We test this key implication of the theory using bioinformatics analysis (see Fig. S5 for the workflow). To that end we explore potential relationships between catalytic efficiencies of enzymes and the length of reaction pathways (Fig. 4D and S6) to which they belong. We represent the metabolic pathway length for each enzyme by the average number of enzymes in its associated pathways. Interestingly, we see significant differences (p < 0.001) of metabolic pathway length distribution between diffusion-limited (red box in Fig. 4D, *k*_*cat*_/*K*_*M*_ > 10^7^ *M*^−1^ *s*^−1^) and reaction-limited (blue box in Fig. 4D, *k*_*cat*_/*K*_*M*_ < 10^7^ *M*^−1^ *s*^−1^) enzymes. Both the median (Fig. 4D) and mean (Fig. S6) metabolic pathway length of diffusion-limited enzymes is smaller than those of reaction-limited enzymes. It is pertinent to highlight that the pathway lengths under consideration significantly exceeds the conventional lengths of metabolic reaction pathways; for instance, the purine biosynthesis pathway contains only ten enzymes ^4, 20^. This discrepancy arises from the inclusion in the KEGG pathway database of all possible branches, connections, and verified reactions in metabolic pathways. Consequently, even rarely occurring or condition-specific reactions contribute to the pathway’s defintion in KEGG. Furthermore, the co-occurrence of diffusion-limited and reaction-limited enzymes within extensive metabolic pathways adds noise into the metabolic pathway length distribution, obfuscating their separation.

## Discussion

Spatiotemporal organization of biomacromolecules and ligands in the cell plays a critical role in modulating biochemical activity. In the past decade, several reversible biomolecular compartments, also known as membraneless organelles (MLOs) have been identified in both prokaryotic and eukaryotic systems. Among several implicated functions of these MLOs is their potential role as biochemical crucibles which promote certain biochemical reactions by colocalizing enzymes and substrates. Indeed, co-localizing enzymes and substrates has been observed to improve reaction efficiencies by several orders of magnitude in engineered biochemical systems ^19^. Further evidence for a potential biological role of clustered enzymes comes in the observation of purinosomes, a reversible, multi-enzyme compartment that co- localizes enzymes of the purine biosynthesis pathway ^4, 20^. Among the potential functions of such a dynamic multi-enzyme complexes is the increase of metabolic flux by improving the efficiency of intermediate processing.

Understanding the mechanistic underpinnings of clustering-induced efficiency of multi-enzyme pathways is therefore crucial not only to obtain insights into the regulation of biochemical pathways but also to design synthetically engineered enzyme droplets^19^. The seminal theoretical work by Castellana et al. ^6^ provides a basis for understanding the speedup of metabolic fluxes in co-clustered enzymatic systems.. However, there is still a need in a comprehensive mechanistic study of the effect of clustering on reaction flixes in enzymatic pathways. Complete understanding of mechanisms modulating enzymatic flux in clustered pathways should reveal the role of different factors, including, crucially the pathway length, strength of protein-protein interactions leading to clustering, reaction rates of individual enzymes and protein-ligand interactions.

Here we we used a combination of approaches to systematically investigate the mechanism by which enzyme clustering speeds up the rate of the final product in linear enzymatic pathways. Our key finding is that clustering is beneficial only for less efficient, reaction-limited enzymes which require several unproductive collisions before the reaction can occur. Crucially, according to detailed studies on kinetic efficiencies of enzymes by Bareven et al. ^11^, 98% or more enzymes are at least one order slower than the diffusion-limit, with the average enzyme being at least 5 orders of magnitude less efficient. This means that only one in 10^4^-10^5^ enzyme-substrate encounters is productive. These numbers are consistent with the kinetic regime wherein we observe the highest speedup in reaction flux (vs the bulk) upon clustering of enzymes (Fig. 4C). On the other hand, the “perfect” diffusion-limited enzymes which comprise < 2% of all enzymes by Bar-even et al. ^11^ show no gain in efficiency upon clustering (Fig.4C). Therefore, clustering of enzymes could improve reaction efficiencies for most natural enzymes by several orders of magnitude over that of the bulk.

Our theory predicts that reaction-limited enzymes would gain most from clustering when they populate long metabolic pathways. We posited therefore that longer pathways should be enriched in less efficient reaction-controlled enzymes while diffusion-limited enzymes should populate shorter pathways. This is somewhat counterintuitive as one might expect that longer pathways require more efficient enzymes to function efficiently in multi-step reactions. We tested this key implication from theory using bioinformatics analysis (Fig. 4D). Crucially, we found a pronounced disparity in terms of pathway lengths between diffusion-limited and reaction-limited enzymes, such that more efficient enzymes populate shorter pathways. It is intriguing to infer that enzyme clustering represents an evolutionary strategy aimed at accomplishing dual objectives: leveraging the same enzymes to catalyze multiple reactions (and therefore delivering lower catalytic efficiency for each reaction) and counterbalancing the ensuing loss of efficiency by enzyme clustering. More generally it appears that spatial clustering of enzymes belonging to the same pathway via weak interactions (2 kT in Fig. 1 and 4) could be a broad evolutionary mechanism to speed up the reaction fluxes in linear enzymatic pathways than evolving highly efficient enzymes which specialize in processing a specific substrate.

Cells could therefore exploit “imperfect”, promiscuous enzymes which catalyze several different reactions in a sub-optimal manner with the tradeoff of requiring several futile encounters for a productive collision ^11^. Our results suggest that the sub-optimal nature of the microscopic properties which characterize the catalytic efficiencies of these enzymes can be overcome by co-clustering the components together providing a balance of efficiency and potential evolvability of new functions and specificities. These results (Fig. 1 and 4) show that the increased local density of enzymes within the cluster coupled with the presence of several copies of enzymes within the cluster can result in faster processing of intermediates along the pathway. Therefore, even when a substrate/intermediate binds to a non-specific protein surface or to an incorrect enzyme and diffuses away, there exist other copies of the cognate enzyme in the immediate vicinity within the cluster which could result in a more efficient turnover. This is further optimized by the presence of weak non-specific attractive interactions between enzymes and substrates (Fig. 3 and S3), allowing them to dwell in the cluster long enough for a reaction to occur before diffusing away. These generic physical mechanisms (Fig. 5) can thus improve reaction efficiencies without actually fine tuning the microscopic properties of the enzyme or its biochemical specificity to a substrate. Evolution could, therefore, use the density-modulating role of enzyme clusters to achieve the twin goals of using the same enzymes to catalyze several reactions while also offsetting the tradeoff in efficiency that accompanies promiscuity.

**Figure 5.**
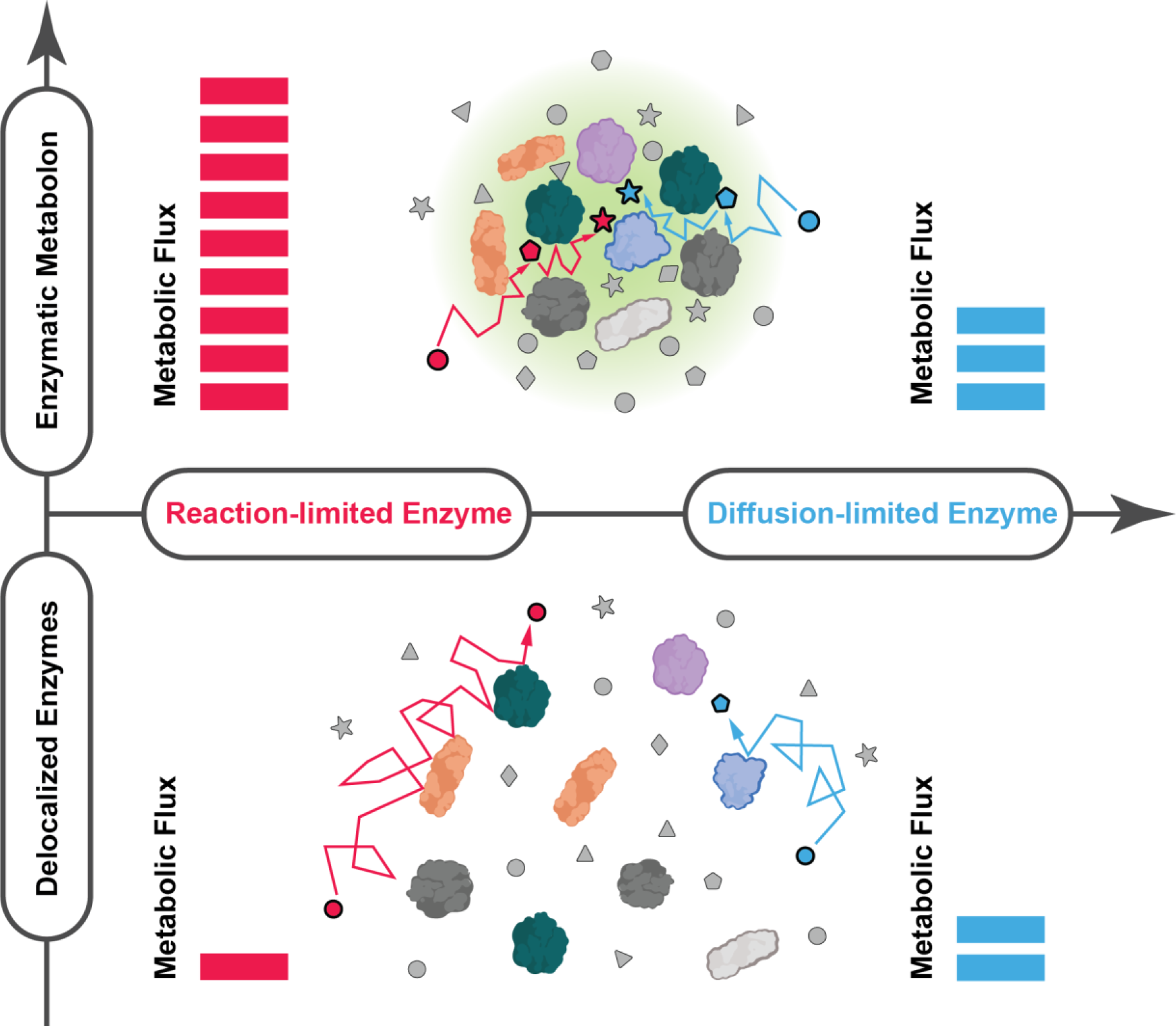
Graphical Summary. Facilitated by non-specific protein-protein interactions, enzymatic metabolon (upper panel) accelerates metabolic flux as contrasted with delocalized enzymes (lower panel). This acceleration is particularly notable in long reaction pathways. Moreover, the impact of this metabolic flux acceleration is markedly pronounced for reaction-limited enzymes (left panel) compared to diffusion-limited enzymes (right panel). This could be attributed to the substantial reduction in the first-passage time facilitated by enzyme clustering in reaction-limited scenarios.

