## Supplementary Information for "Enzymatic metabolon improves kinetic efficiency of reaction-limited enzyme pathways"

Srivastav Ranganathan[#], Junlang Liu[#], and Eugene Shakhnovich[*]
Department of Chemistry and Chemical Biology, Harvard University, Cambridge, MA 02138

[#] These authors contributed equally to this work.

**This PDF file includes:**
Methods
Figs. S1 to S7
References

**Other supplementary material for this manuscript includes the following:**
Supplementary Form

### Methods

**Mathematical Derivation of Reaction Efficiency**

Equation S1 (Eq. 9 in main text) gives the reaction efficiency $\eta$ of delocalized scenario where both enzymes and substrates are uniformly distributed in the reaction unit. It can be expressed as follows,

$$\eta_{\text{delocalized}} = \frac{(k\cdot n)^l}{(k\cdot n+\alpha_0)(k\cdot n+\beta)^{l-1}}, \tag{S1}$$

where $k$ is the catalytic efficiency for all reactions in the pathway, $n$ is the density of any enzyme in the pathway based on the assumption that all enzymes in the pathway have the same density and operate at the same catalytic efficiency.

Here is the mathematical derivation of Equation S1. Since 1) both enzymes and substrates are uniformly distributed in the reaction unit; 2) all enzymes in the pathway have the same density and operate at the same catalytic efficiency, reaction-diffusion equation (Eq. 4-5 in main text) for delocalized scenario can be simplified as

$$\partial_{\mathrm{t}} c_0(\mathrm{t}) = \; -k\cdot n\cdot c_0(\mathrm{t}) - \alpha_0(c_0(\mathrm{t}) - c_0^*), \tag{S2}$$

$$\partial_{\mathrm{t}} c_i(\mathrm{t}) = \; k\cdot n\cdot c_{i-1}(\mathrm{t}) - k\cdot n\cdot c_i(\mathrm{t}) - \beta\cdot c_i(\mathrm{t}), \tag{S3}$$

where $i$ ranges from 1 to $l$ for a enzymatic reaction pathway with $l$ chain reactions. At steady state, we have

$$\partial_{\mathrm{t}} c_0(\mathrm{t}) = \; 0, \tag{S4}$$

$$\partial_{\mathrm{t}} c_i(\mathrm{t}) = \; 0. \tag{S5}$$

From Eq. S3, we have steady-state concentration of substrate $S_0$ as

$$c_0 = \; \frac{\alpha_0\cdot c_0^*}{k\cdot n + \alpha_0}\,. \tag{S6}$$

From Eq. S4 and S5, we have steady-state concentration of substrate $S_i$ (for $i = 1 \ldots l$) as

$$c_i = \; \frac{k\cdot n}{k\cdot n + \beta}\cdot c_{i-1} = \frac{\alpha_0\cdot c_0^*\cdot (k\cdot n)^i}{(k\cdot n+\alpha_0)(k\cdot n+\beta)^i}\,. \tag{S7}$$

Therefore, we can express the reaction efficiency $\eta$ of delocalized scenario as follows according to Eq. S8,

$$\eta_{\text{delocalized}} = \frac{k_l \int_0^R dr\;\, 4\pi r^2 n_l(r) c_{l-1}(r)}{\alpha_0 (4/3)\pi R^3 c_0^*} = \frac{k\cdot n\cdot c_{l-1}\int_0^R dr\; 4\pi r^2}{\alpha_0 (4/3)\pi R^3 c_0^*} = \frac{(k\cdot n)^l}{(k\cdot n+\alpha_0)(k\cdot n+\beta)^{l-1}}, \tag{S8}$$

where the rightmost part is Eq. S1.

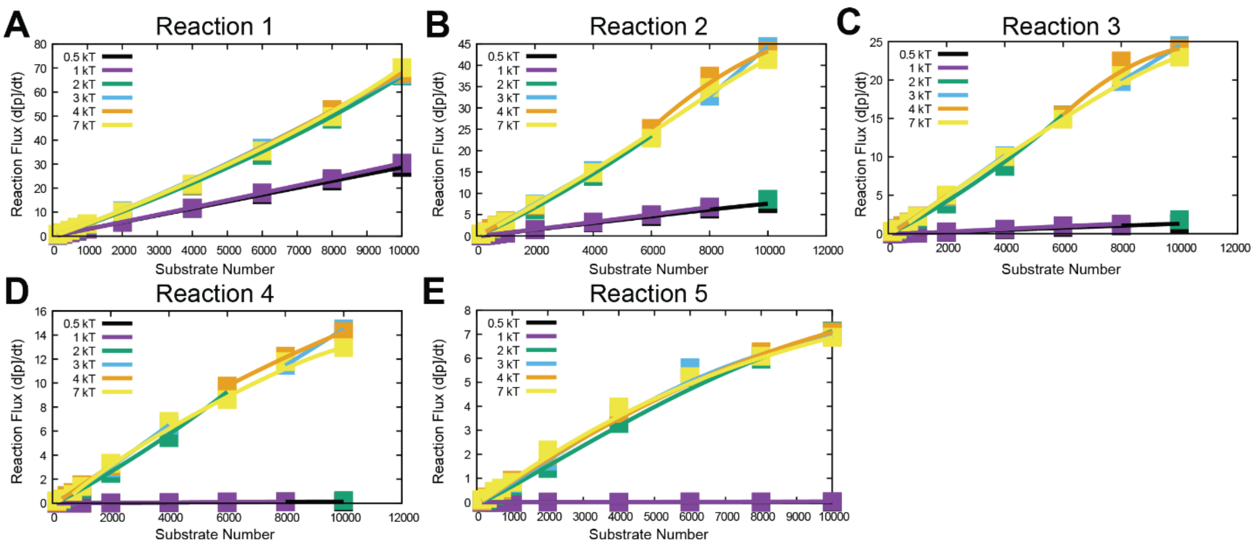


**Figure S1. Reaction flux vs. substrate concentration curves for different reactions in a 5-reaction metabolic pathway.** A)-E) show reaction flux vs. substrate concentration curves for the first (A), second (B), third (C), fourth (D) and fifth (E) reaction respectively. Different colors represent different protein-protein interaction strengths ($\epsilon_{protein-protein}$).

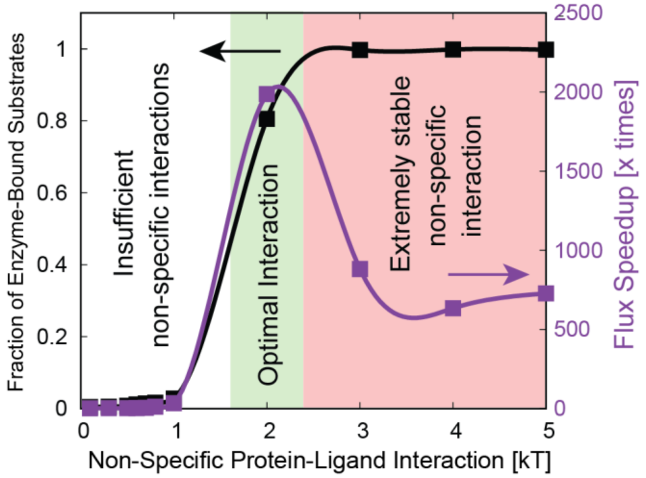


**Figure S2. Importance of non-specific protein-ligand interactions.** Speedup in reaction flux shows a dramatic increase upon introduction of non-specific protein-ligand interactions. A further increase in non-specific protein-ligand interactions results in a dropoff in flux (Purple Curve). The black curve shows the fraction of ligand molecules bound to the enzymes.

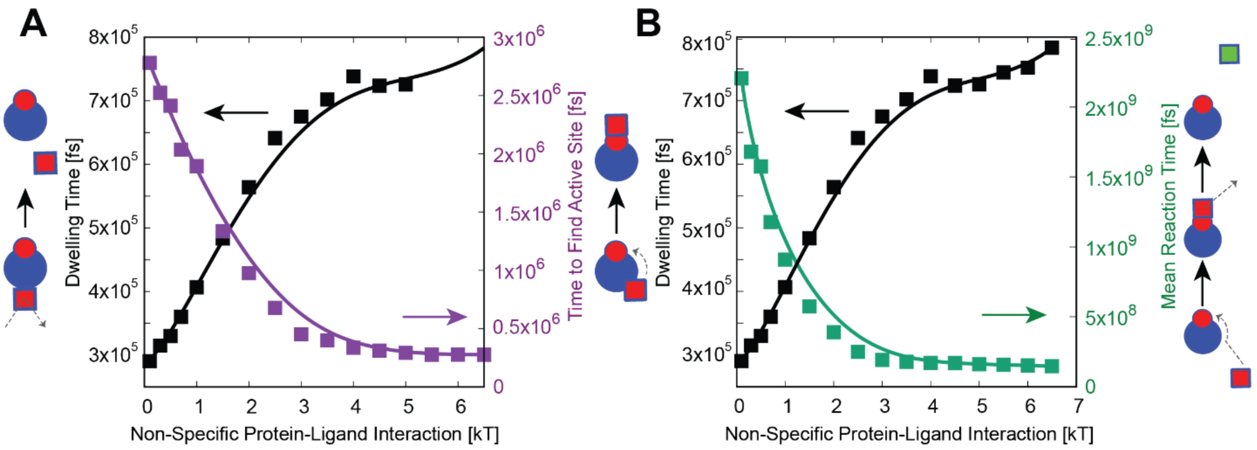


**Figure S3. Changes of different timescales as a function of non-specific protein-ligand interactions in two-particle (enzyme-substrate) system.** A) Black Curve - The mean dwelling time, the time spent by the ligand in contact with the enzyme before it diffuses away to solution ($T_{dwell}$) in two particle enzyme-substrate simulations. Purple Curve - The mean first passage time for the ligand to find the binding site upon coming in contact with the enzyme ($T_{find}$) in two particle enzyme-substrate simulations. An increase in non-specific interaction strength results in an increase in the dwelling time of the ligand on the enzyme surface while concomitantly resulting in a speedup in the search process of the binding site. B) Black Curve – Same as Fig. S5A. Green Curve- Mean first passage time to go from the free state (enzyme and substrate unbound) to the formation of a product ($T_{react}$).

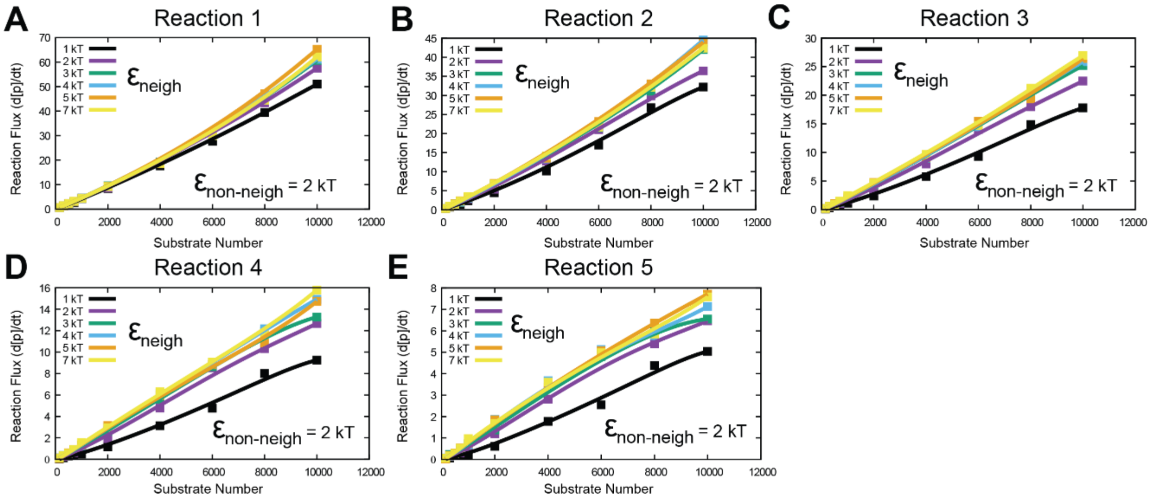


**Figure S4. Spatial seggregation of enzymes within the enzyme cluster offers only marginal speedup in reaction fluxes.** Reaction flux vs. substrate concentration curves for a 5-reaction linear metabolic pathway. A)-E) correspond to the first (A), second (B), third (C), fourth (D) and fifth (E) reaction respectively. Here, neighboring enzymes (such as, enzymes for the first and second reactions) in the pathway interact via a different interaction strength compared to non-neighbor enzymes (such as, enzymes for the first and third/forth/fifth reactions). In our simulations so far, we consider all protein-protein interactions to be homogenous and isotropic. In other words, there is a uniform spatial distribution of enzymes within the cluster. However, biological interactions could display heterogeneity. Further, tunneling of substrate from one enzyme to another could also result in a gain in specificity and efficiency of enzyme clusters. In this context, we look at a scenario where successive enzymes in the pathway interact at different interaction strengths $\epsilon_{neigh}$ while all other pairwise protein-protein interactions $\epsilon_{non-neigh}$ are set to 2 kT, a regime where clusters are favored (Fig. 1C). In Fig. S2A-E, we systematically vary $\epsilon_{neigh}$ from 1 kT ($\epsilon_{neigh} < \epsilon_{non-neigh}$) to 7 kT. Across all the 5 reactions, the biggest speedup in velocity was observed as we switch from $\epsilon_{neigh}$ of 1 kT ($\epsilon_{neigh} < \epsilon_{non-neigh}$) to 2 kT ($\epsilon_{neigh} = \epsilon_{non-neigh}$). A further increase in $\epsilon_{neigh}$ so as to favor interactions between successive reactions only results in a marginal increase in reaction flux. Unlike the gain in reaction flux upon formation of enzyme clusters (Fig. 1C, $\epsilon_{protein-protein} \geq$ 2 kT) and the introduction of weak non-specific protein-ligand interactions (Fig. 3), there is no significant gain evident from introducing favorable interactions between successive enzymes of the pathway. Therefore, weak isotropic non-specific interactions can sufficiently accelerate long reaction pathways even in the absence of specific protein-protein interaction networks.

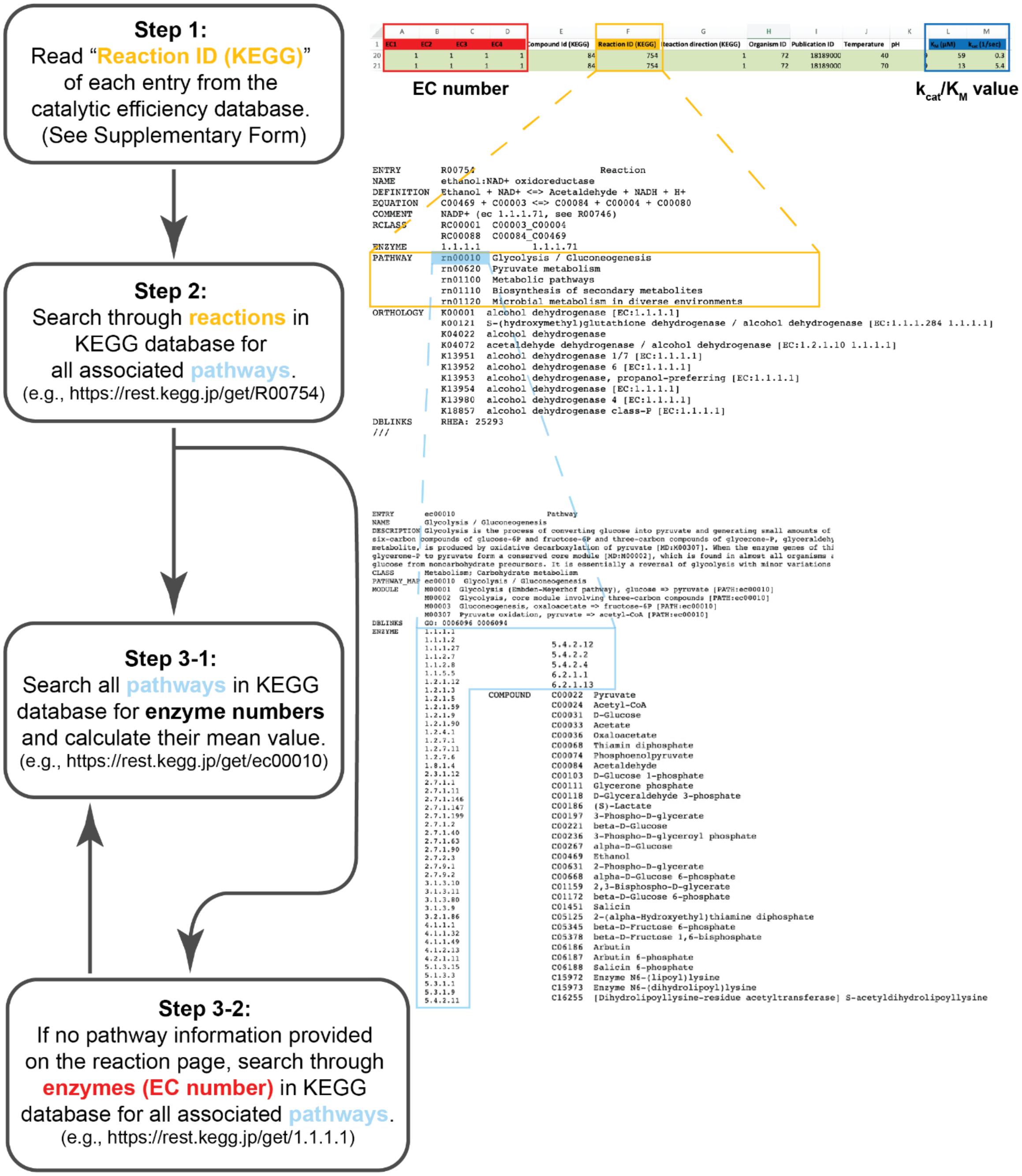


**Figure S5. Workflow of the bioinformatics study used in this work.** We obtained 4550 catalytic efficiency entries from the database Ref. 1. After analysis following above workflow, we obtained 4239 catalytic efficiency entries with average enzyme number in their associated pathways using KEGG database [2], among which there are 446 diffusion-limited ($k_{cat}/K_M > 10^7\ M^{-1}\,s^{-1}$) enzyme entries and 3793 reaction-limited ($k_{cat}/K_M < 10^7\ M^{-1}\,s^{-1}$) enzyme entries.

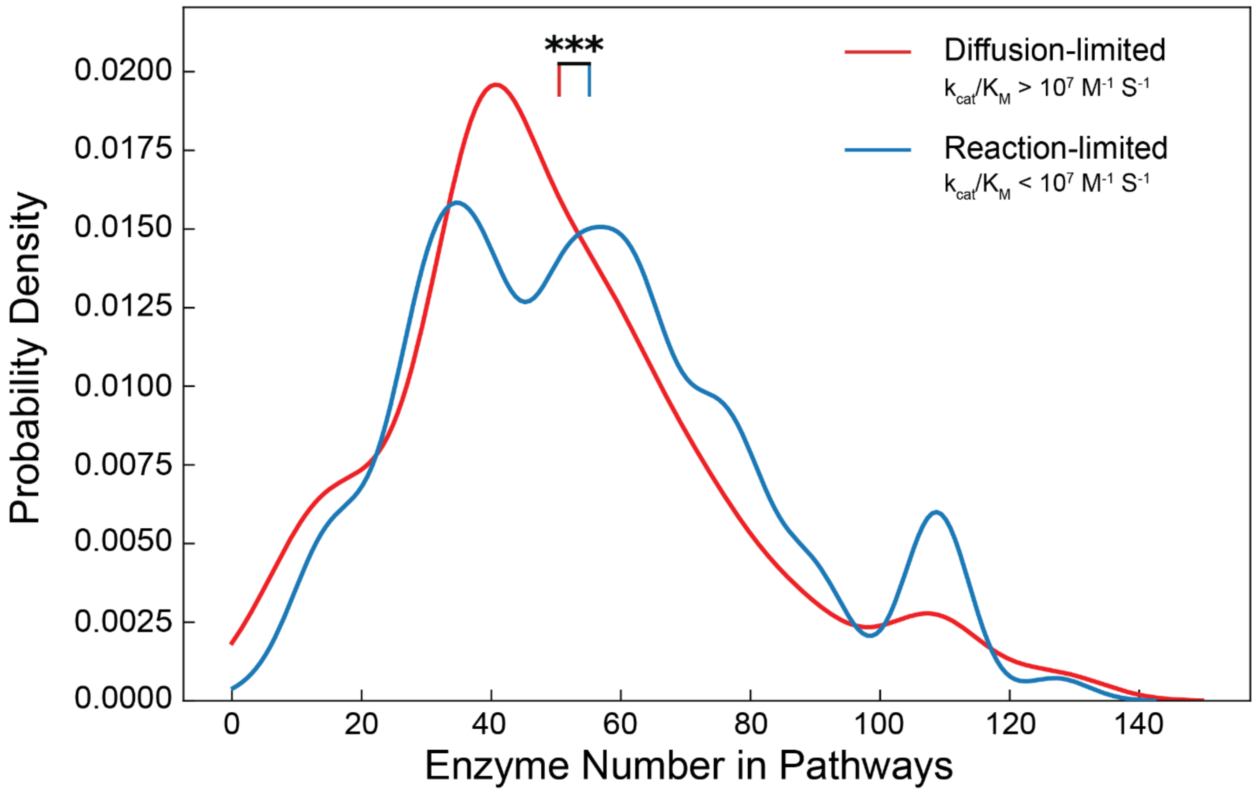


**Figure S6. Distribution of average enzyme number in associated pathways for both diffusion-limited and reaction-limited enzymes.** 3-star (***) means p-value is less than 0.001. The corresponding mean values of each distribution are denoted by diminutive red and blue vertical bars positioned beneath the 3-star.
